# Anti-diarrheal drug loperamide induces dysbiosis in zebrafish microbiota via bacterial inhibition

**DOI:** 10.1101/2023.04.18.537295

**Authors:** Rebecca J. Stevick, Sébastien Bedu, Nicolas Dray, Jean-Marc Ghigo, David Pérez-Pascual

## Abstract

**Background:** Perturbations of animal-associated microbiomes from chemical stress can affect host physiology and health. While dysbiosis induced by antibiotic treatments and disease are well known, chemical, non-antibiotic drugs have recently been shown to induce changes in microbiome composition, warranting further exploration. Loperamide is an opioid-receptor agonist drug widely prescribed drug for treating acute diarrhea in humans. Loperamide is also used as a tool to study the impact of bowel dysfunction in animal models by inducing constipation, but its effect on host-associated microbiota is poorly characterized.

**Results:** We used conventional and gnotobiotic larval zebrafish models to show that in addition to host-specific effects, loperamide also has anti-bacterial activities that directly induce changes in microbiota diversity. This dysbiosis is due to changes in bacterial colonization, since germ-free zebrafish mono-reconventionalized with bacterial strains sensitive to loperamide are colonized up to 100-fold lower when treated with loperamide. Consistently, the bacterial diversity of gnotobiotic zebrafish colonized by a mix of representative bacterial strains is affected by loperamide treatment.

**Conclusion:** Our results demonstrate that loperamide, in addition to host effects, also induces dysbiosis in a vertebrate model, highlighting that established treatments can have underlooked secondary effects on microbiota structure and function. This study further provides a insights for future studies exploring how common medications directly induce changes in host-associated microbiota.

**Graphical Abstract:** 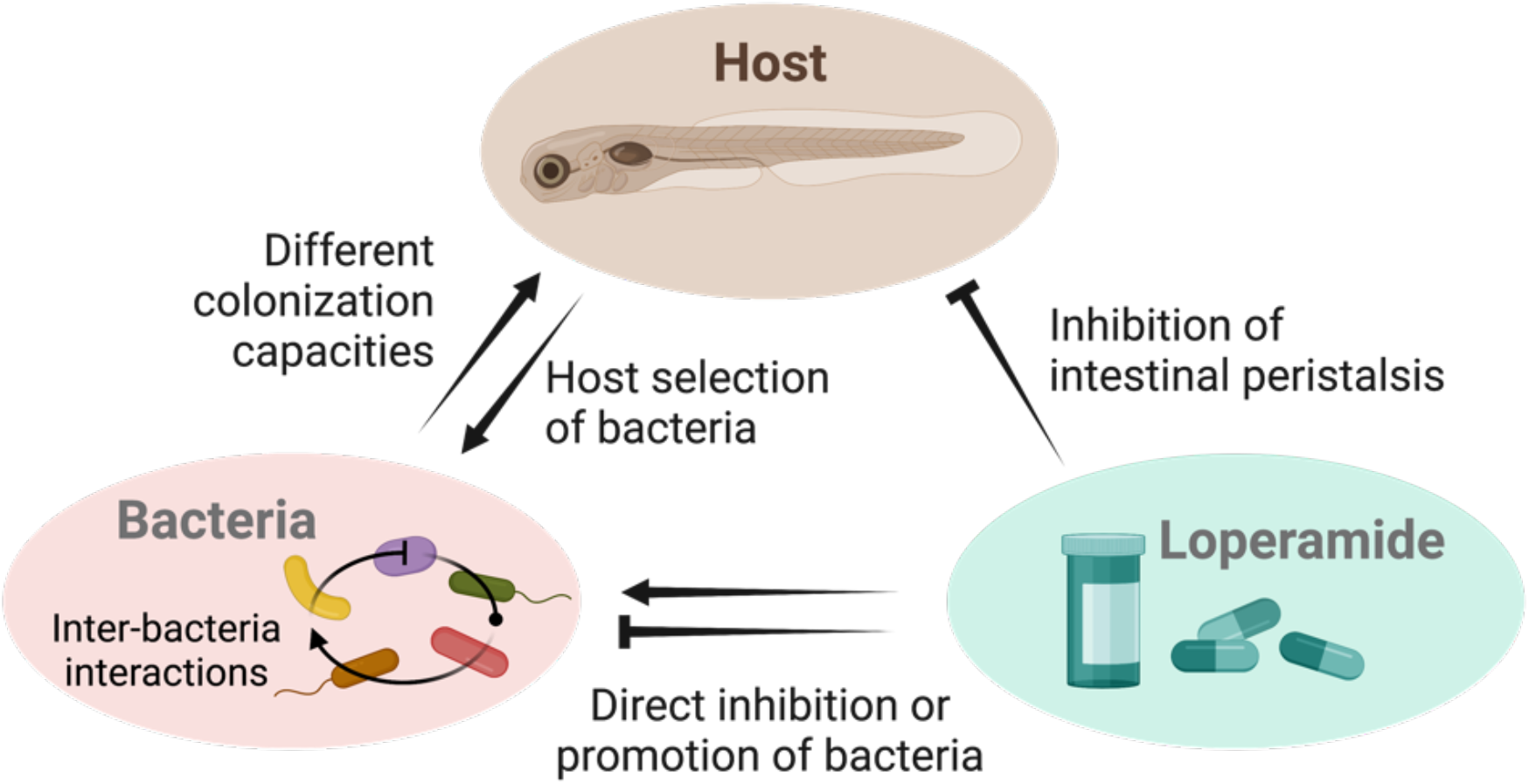

## Background

Animal-associated microbiomes are dynamic communities that play essential roles in the physiology, health, and evolution of their hosts [1]. Numerous studies have explored the impact of different phenomena on microbiota stability, including antibiotic treatments, gut health, or environmental factors, in order to understand the consequences of microbiota perturbations on host functions [2–5]. These perturbations may lead to dysbiosis or a change in microbial community composition and/or function, relative to the steady state, with potential implications for host health [6, 7].

While the complexity of microbiota in humans and animal models limits functional and mechanistic studies, germ-free and gnotobiotic animal models with controlled, tractable microbiota are widely used to study host-microbiota interactions [8]. Compared to conventional animals with relatively variable microbiota [9], gnotobiotic animals with host-specific bacterial consortia can mimic key phenotypes for mechanistic studies, and are powerful tools to simplify microbiota and increase experimental reproducibility [10]. In particular, zebrafish (*Danio rerio*), which possesses both an innate and adaptative immune system and a mammal-like intestinal epithelium, has emerged as an established gnotobiotic model to study vertebrate host-microbiota interactions [11, 12]. Gnotobiotic larval zebrafish can indeed be easily reared to study simplified host-microbial systems in the context of developmental biology, immunology, and disease [13, 14].

Loperamide is a prevalent medication for treating diarrhea in humans and animals that acts on µ-opioid receptors in the large intestine, decreasing intestinal peristaltic activity and increasing the absorption of fluids [15–18]. Loperamide is also used to study bowel dysfunction and constipation in animal models, including rats, mice, and zebrafish, generating a relevant model of irritable bowel syndrome or opioid-induced bowel dysfunction disorder [19–21]. In zebrafish, loperamide treatment was shown to cause a significant decrease in intestinal peristaltic frequency that can be restored by the presence of specific bacteria or acetylcholine [21, 22].

Despite its pervasive use in humans and animal models, the potential effects of loperamide on host-associated microbiota *in vitro* and *in vivo* are poorly characterized. It has been suggested that slow transit time and constipation induced by loperamide could be responsible for changes in bacterial composition and decreased diversity observed in rats and mice [23–26]. Although publications using loperamide to investigate host-associated microbiota establish the link between constipation and microbial dysbiosis, recent studies have identified loperamide hydrochloride and its derivatives as molecules displaying bactericidal activity [27–29]. Hence the extent of microbial dysbiosis directly caused by this compound versus its impact on the alteration of host function are poorly understood and ignored in animal models.

In this study, we used conventional and gnotobiotic larval zebrafish to reproduce *in vivo* loperamide-induced dysbiosis based on *in vitro* bacterial sensitivity to loperamide. We found that loperamide leads to recoverable dysbiosis in conventional larval zebrafish according to strain-specific inhibition or promotion of bacteria. Our results demonstrate how a relevant chemical perturbation induces dysbiosis in a vertebrate microbiome model. These findings should be considered in the context of secondary effects of established treatments, assumed mode of action in animal models, and microbiota community recovery.

## Methods

### General zebrafish husbandry

Wild-type AB/AB zebrafish (*Danio rerio*) fertilized eggs at 0 days post fertilization (dpf) were obtained from the Zorgl’hub platform at Institut Pasteur. All procedures were performed at 28°C under a laminar microbiological hood with single-use disposable plastic ware according to European Union guidelines for handling of laboratory animals and were approved by the relevant institutional Animal Health and Care Committees. Eggs were kept in 25 cm^3^ vented flasks (Corning 430639) with 20 mL of autoclaved mineral water (Volvic) until 4 dpf (30 – 33 eggs/flask) and transferred to new flasks after hatching at 4 dpf (10 – 15 fish/flask). At 6 dpf, each fish was transferred to an individual well of a 24-well plate (TPP 92024) in 2 mL of autoclaved mineral water and maintained until the end of the experiment (11 dpf). Conventional zebrafish embryos were transferred to flasks at 1 dpf and maintained as described. At the end of the experiment, zebrafish were euthanized with an overdose of tricaine (MS-222, Sigma-Aldrich E10521) at 0.3 mg/mL for 10 minutes.

Fish were fed with sterile *Tetrahymena thermophila* every 48 hours starting at 4 dpf. Germ-free *T. thermophila* stocks were kept in 15 mL of PPYE broth (0.25% protease peptone BD Bacto #211684, 0.25% yeast extract BD Bacto #212750) supplemented with penicillin G (10 unit/mL) and streptomycin (10 µg/mL) at 28°C. Every week, a new stock was inoculated with 100 µL of the previous stock and tested for sterility on LB, TYES, and YPD agar media plates. To prepare food for the zebrafish, *T. thermophila* was inoculated at a 1:50 ratio from the stock into 20 mL MYE broth (1% milk powder, 1% yeast extract) and grown for 2 days. On feeding day, the *T. thermophila* was transferred to a 50 mL Falcon tube and washed 3 times (4400 rpm, 3 min at 25 °C) with sterile mineral water. Resuspended *T. thermophila* was added to the fish in culture flasks (500 µL in 20 mL) or 24-well plates (50 µL in 2 mL).

### Germ-free zebrafish sterilization

The zebrafish embryos were sterilized as previously described with the following modifications [13, 30]. Recently fertilized zebrafish eggs (0 dpf) were bleached (0.000005 % final v/v) for 5 minutes, then washed 2 times in sterile mineral water. Eggs were then maintained in 50 mL Falcon tubes (100 eggs/tube) overnight in 35 mL of sterile mineral water supplemented with 0.4 µg/mL methylene blue solution (Sigma Aldrich 50484). At 1 dpf, the volume of each tube was adjusted to 50 mL and the eggs were treated with an antibiotic cocktail for 2 hours with gentle agitation at 10 rev/min: penicillin G: streptomycin at 100 µg/mL (GIBCO 15140148), kanamycin sulfate at 400 µg/mL (PAN BIOTECH P06-04010P) and amphotericin B solution at 250 µg/mL (Sigma-Aldrich A2942). Then, the eggs were washed 3 times with sterile mineral water and resuspended in 50 mL water. The eggs were bleached (0.000005 % final v/v) for 15 minutes with inversion every 3 minutes, then washed 3 times in sterile mineral water and resuspended in 50 mL water. Finally, the eggs were treated with 1 % Romeoid solution (COFA, France) for 10 minutes, then washed 3 times in sterile mineral water. Eggs were then transferred to 25 cm^3^ vented flasks and maintained as described above.

Sterility was confirmed at 3 dpf by spotting 50 µL of water from each flask on LB, TYES and YPD agar plates and incubated at 28 °C under aerobic conditions for at least 3 days. In addition, monthly checks of bacterial contamination were done by PCR amplification of water samples with 16S rRNA gene primers as described below in the characterization section. Contaminated flasks were immediately removed from the experiment and not included in the results.

### Germ-free zebrafish re-conventionalization

Germ-free zebrafish larvae were re-conventionalized at 4 dpf, as follows. Overnight cultures of a single bacterial colony in 5 mL of liquid media were washed twice with sterile Volvic water and normalized to OD-0.1 in water. For mono-reconventionalization, 200 µL of bacterial suspension was added into flasks of germ-free zebrafish in 20 mL of Volvic water at a final concentration of 5 × 10^5^ CFU/mL. For mix-reconventionalization, 200 µL of each strain was added per flask at a final concentration of 5 × 10^5^ CFU/mL per strain. Water samples were plated in serial dilutions to confirm final bacterial concentration and sterility. Re-conventionalization was performed for 48 hours until fish were transferred to sterile water in 24-well plates.

### Zebrafish loperamide treatment

Loperamide hydrochloride (Sigma-Aldrich 34014) was dissolved in pure dimethyl sulfoxide (DMSO, Sigma-Aldrich D8418) at a stock concentration of 100 mg/mL. Larval zebrafish were treated at 5 dpf with loperamide at a final concentration of 10 mg/L in 20 mL vented flasks for 24 hours, which has been previously shown to significantly reduce peristaltic movement in larval zebrafish at 4 - 6 dpf [21]. Sterile DMSO added at a final concentration of 1:10000 was used as the control. After 24 hours of treatment, all 6 dpf fish were transferred to water and maintained until sampling.

### Conventional zebrafish sampling and DNA extraction

Zebrafish larvae were sampled at each of 3 timepoints (6 dpf, 7 dpf, 11 dpf) with 3 treatment conditions (control water, DMSO 1:10000, Loperamide 10 mg/L) as follows. At each timepoint, 5 larval fish per condition (15 total) were washed twice by transfers to clean, sterile water to remove environmental and residual bacteria. Each fish was then added to a sterile 2-mL microcentrifuge tube in 200 µL of water and euthanized with tricaine at 0.3 mg/mL. All liquid was removed from the tissue, then the samples (45 total) were immediately frozen at - 80°C and stored until DNA extraction.

DNA extraction was performed from single larval zebrafish using the DNeasy Blood & Tissue kit (Qiagen 69504) with modifications as follows. Tissue samples were thawed at room temperature, then 380 µL Buffer ATL and 20 µL proteinase K were added directly to each individual larva in a 2 mL tube. Samples were vortexed for 15 seconds, then incubated overnight (15-18 hours) at 56°C and 300 rpm until fully lysed. After lysis, 4 µL of RNAse A solution was added and the samples were incubated for 5 minutes at room temperature to remove residual RNA. Next, 400 µL Buffer AL and 400 µL 100 % ethanol were added and mixed by vortexing before loading the lysate onto the DNeasy mini spin column in 2 × 600 µL loads. DNA purification and cleanup proceeded according to the manufacturer’s recommendations with a final elution volume of 50 µL in Buffer AE. Purified DNA was quantified using the Qubit HS DNA fluorometer kit (ThermoFisher Q32851) and purity was assessed with the Nanodrop spectrophotometer (ThermoFisher). DNA yields per single fish sample ranged from 10-15 ng/µL in 50 µL with purity ratios >1.8. Negative controls for the extraction kit were prepared alongside zebrafish samples, but with no tissue input.

### Conventional fish 16S rRNA gene amplicon sequencing and analysis

16S rRNA gene amplicons of the V6 region for the 45 conventional zebrafish samples, 2 mock community samples (Zymo Research DNA standard I D6305), 2 negative DNA extraction samples, and blank PCR control were prepared using 967F/1064R primers. The DNA extraction negative control samples were pooled and concentrated prior to PCR to obtain enough product for sequencing. A two-step PCR reaction using 200 ng of zebrafish DNA was performed in duplicate 50 µL reactions as previously described [31, 32]. Each first step reaction included 25 µL 2X Phusion Mastermix (Thermo Scientific F531S), 1.5 µL of 10 µM F/R primer mix (967F: CTAACCGANGAACCTYACC, CNACGCGAAGAACCTTANC, CAACGCGMARAACCTTACC, ATACGCGARGAACCTTACC (equimolar mix) / 1064R: CGACRRCCATGCANCACCT), 13 - 20 µL template DNA (200 ng), and 3.5 – 10.5 µL nuclease-free water (up to 50 µL). PCR amplification (step 1) conditions were denaturing at 98°C for 3 min followed by 30 cycles of denaturation at 98°C for 10 s, primer annealing at 56°C for 30 s, and extension at 72°C for 20 s, then a final extension at 72°C for 20 s. Negative controls for the PCR reagents were prepared alongside zebrafish DNA samples, but with additional nuclease-free water input. PCR products were assed for concentration (Qubit DNA HS reagents) and expected size using agarose gel electrophoresis. A second PCR step was performed to attach sequencing barcodes and adaptors according to Illumina protocols. The PCR products were analyzed with 250 bp paired-end sequencing to obtain overlapping reads on an Illumina MiSeq at the Institut Pasteur Biomics platform.

The resulting 16S rRNA gene amplicon sequences were demultiplexed and quality filtered using DADA2 (v1.6.0) implemented in QIIME2 (v2020.11.1) with additional parameters --p-trunc-len-r 80 --p-trunc-len-f 80 --p-trim-left-r 19 --p-trim-left-f 19 to determine amplicon sequence variants (ASVs) [33, 34]. All ASVs were summarized with the QIIME2 pipeline (v2020.11.1) and classified directly using the SILVA database (99 % similarity, release #134) [35, 36]. Processed ASV and associated taxonomy data was exported as a count matrix for analysis in R (v4.1.3). The positive and negative controls were checked to ensure sequencing quality and expected relative abundances. Non-bacterial and chloroplast sequences were then removed, and the data was normalized by percentage to the total ASVs in each sample for further dissimilarity metric analysis.

All descriptive and statistical analyses were performed in the R statistical computing environment with the *tidyverse* v1.3.1, *vegan* v2.5.7 and *phyloseq* v1.38.0 packages [37–39]. Rarefaction curves and sequencing coverage estimates were generated using the rarecurve() commands with sample=[number of reads in smallest sample] in *vegan* v2.5.7 [40]. Non-metric dimensional analysis (NMDS) was used to determine the influence of timepoint or loperamide treatment on the ASV-level composition. The Bray-Curtis dissimilarity metric was calculated with k = 2 for max 50 iterations and 95 % confidence intervals (standard deviation) were plotted. Statistical testing of the beta-diversity was done using the PERMANOVA *adonis2* test implemented in *vegan* (method = “bray”, k = 2) [41, 42]. Within-condition variability was calculated using the command vegdist(method = “bray”, k = 2) and the matrix was simplified to include samples compared within each timepoint.

Significant differences in genera between DMSO (reference) and loperamide-treated (test) at each timepoint were calculated using *limma* implemented in the *microbiomeMarker* v1.1.2 package using the following conditions: norm = “RLE”, pvalue_cutoff = 0.05, taxa_rank = “Genus”, p_adjust = “fdr” [43–45]. Simpson’s diversity values were calculated for each sample at the ASV level using the *vegan* package and analyzed using the non-parametric Kruskal– Wallis rank sum test in R. Additional visualizations were computed using the *ComplexHeatmap* v2.10.0 and *UpSetR* v1.4.0 packages [46, 47]. All processed sequencing files, bash scripts, QIIME2 artifacts, and Rmd scripts to reproduce the figures in the manuscript are available on Zenodo [48].

### Measurement of zebrafish growth and development

In order to determine the effect of loperamide growth on larval fish growth and development, 9-10 fish were sampled at each timepoint (6, 7, 11 dpf) for each condition (control water, DMSO, loperamide) = 85 fish total. After euthanasia, the samples were fixed in 1 % paraformaldehyde (PFA) and stored at 4°C. After fixation, the samples were rinsed 3 times with PBS then placed into individual wells in a plate. Microscopy images were taken with a Leica M80 10X with a Leica IC80 HD camera. Four images were captured per sample: whole at 2.5X, caudal at 5X, lateral at 5X and head at 5X for a total of 337 images for 85 samples. Relevant measurements of each fish sample were performed using ImageJ [49]. Four measurements in millimeters were taken per fish: eye diameter, rump-anus length, standard length, and tail width according to methods previously described [50–52].

### Bacterial strains and growth conditions

Bacterial strains are listed in Supplementary Table S1. Zebrafish-associated strains were grown in Tryptone Yeast Extract Salts (TYES) or Miller’s Lysogeny Broth (LB) (Corning) and incubated at 28°C with rotation. Cultures on solid media were on LB or TYES with 1.5 % agar. Bacteria were always streaked from glycerol stocks on LB- or TYES-agar before inoculation with a single colony in liquid cultures. All media and chemicals were purchased from Sigma-Aldrich.

### Isolation and 16S characterization of bacteria from conventional zebrafish

Five of the zebrafish-associated strains were previously isolated and characterized from the zebrafish environment [53]. The following strains were isolated and identified in the same way in this study: S2, S4, S8, S9. Zebrafish lysates and tank water were serially diluted and plated on R2A, TYES, and LB agar and incubated at 28°C for up to 3 days. Each colony morphotype per media was catalogued and re-streaked on the same agar. The morphotype identification was done as previously described [53, 54]. Individual colonies were picked for each morphotype from each agar plates, vortexed in 200 µl DNA-free water and boiled for 10 min at 90°C. Five µl of this bacterial suspension was used as template for colony PCR to amplify the 16S rRNA gene with the universal primer pair 27f and 1492R. 16S rRNA gene PCR products were verified on 1% agarose gels, purified with the QIAquick PCR purification kit (Qiagen) and two PCR products for each morphotype were sent for sequencing (Eurofins, Ebersberg, Germany). Individual 16S rRNA-gene sequences were compared with those available in the EzBioCloud database. Species-level identification was performed based on the 16S rRNA gene sequence similarity was >99%. The zebrafish-associated strains used in this study (Table S1) were chosen from this catalogue based on their sensitivity to loperamide and match with significant changes in the conventional 16S rRNA gene amplicon data.

### Bacterial growth curves and survival assays

Overnight cultures of a single bacterial colony in 5 mL of liquid media were measured and normalized to OD-0.5. Liquid media supplemented with 10 mg/L loperamide in DMSO or 1:10000 DMSO or control was added to a TPP flat-bottom polystyrene 96-well plate. Bacterial cultures were added to each condition in triplicate at a final starting concentration of OD-0.05 in 100 µL. Negative control wells were included for each media and condition. A plastic adhesive film (adhesive sealing sheet, Thermo Scientific, AB0558) was used to seal the wells, and the plates were then incubated in a TECAN Infinite M200 Pro spectrophotometer for 20 hours at 28°C. OD600 was measured every 30 minutes, after a 30-second orbital shaking of 2 mm amplitude.

Bacterial survival in water was tested using the *in vivo* re-conventionalization conditions described above. Overnight cultures of a single bacterial colony in 5 mL of liquid media were washed twice with autoclaved Volvic water, measured and normalized to OD-0.1 in water. Bacteria were inoculated at a final concentration of 5 × 10^5^ CFU/mL into 10 mL of Volvic water supplemented with loperamide in DMSO at 10 mg/L or 1:10000 DMSO or control. Viable colony forming units (CFUs) were counted from each flask at 0, 6, 24, 48, and 72 hours as follows. Three x 200 µL aliquots were sampled and dilutions were made, then 10 µL drops were plated on LB or TYES and grown at 28°C for 2 days. CFUs were then counted for each strain and CFUs/mL were calculated by 1000 µL/mL / 10 µL plated * dilution factor * (average of replicate CFUs per strain). Survival of each strain was repeated at least two independent times.

### Quantification of gnotobiotic zebrafish bacterial load by CFU counts

Zebrafish were sampled at each of 3 timepoints (6 dpf, 7 dpf, 11 dpf) with 3 treatment conditions (control water, DMSO 1:10000, Loperamide 10 mg/L). At each timepoint, 3-4 larval fish per condition were washed twice by 2 transfers to clean, sterile water in petri dishes to remove environmental and residual bacteria. The larvae were then euthanized with tricaine at 0.3 mg/mL and added in 500 µL of sterile water to 2 mL tubes containing 1.4 mm ceramic beads (Fischer Scientific 15555799). Fish were homogenized for 2 × 45 seconds at 6000 rpm using a 24 Touch Homogenizer (Bertin Instruments). These homogenization conditions are sufficient to lyse zebrafish tissue, but not harmful to the bacteria. The lysate was then diluted from 10-100-fold. For the mono-reconventionalized fish, 10 µL drops were plated in triplicate for each dilution on media. After 2 days of incubation at 28°C, CFUs were counted and CFUs per fish were calculated by 500 µL lysate / 10 µL plated * dilution factor * average of replicate CFUs. For the mix-reconventionalized fish, 3 × 100 µL from each dilution was spread on media using sterile glass beads to differentiate the colonies. After 2 days of incubation at 28°C, CFUs were counted for each strain and CFUs per strain per fish were calculated by 500 µL lysate / 100 µL plated * dilution factor * (average of replicate CFUs per strain).

### Statistical Analyses

All plotting and statistical analyses were performed in the R statistical computing environment (4.1.3) using RStudio (v.2022.02.1) with the *tidyverse* v1.3.1, *ggpubr* v0.4.0, *ggtext* v0.1.1 and *patchwork* v1.1.1 packages [39, 55–57]. Non-parametric global Kruskal-Wallis tests and subsequent Wilcoxon pairwise tests were performed to compare loperamide-treated condition to the DMSO control using compare_means() or stat_compare_means() when p<0.05 is significant. For the comparison of zebrafish colonization and water survival, mean CFUs/mL or CFUs/fish of each strain S1 – S10 were calculated for control conditions at 48 h or T0, respectively. The colonization efficiency for each strain was calculated by *Colonization efficiency = mean CFUs per Fish / mean Water CFUs per mL * 100*. The correlation between the variables was fit with geom_smooth(method = “lm”) and the fit was indicated with correlations using stat_cor() and stat_regline_equation(). Hypothetical bacterial composition comparison of mono-reconventionalized fish was calculated by the mean CFUs per fish per strain / the sum of mean CFUs per fish of S1, S3, S5, S6, S7. Bacterial composition comparison of mix-reconventionalized fish was calculated by the mean CFUs of each strain / the total CFUs of all strains in each fish. Simpson’s diversity values were calculated for each mix-reconventionalized fish based on percent abundance per strain using the *vegan* package and analyzed using the non-parametric Kruskal–Wallis rank sum test in R. All raw data and Rmd scripts to reproduce the figures and statistical tests in the manuscript are available on Zenodo [48].

## Results

### Loperamide treatment induces recoverable dysbiosis in conventional larval zebrafish microbiota

Using the experimental procedure described in Figure 1, we determined the impact of loperamide treatment on conventional larval zebrafish microbiota using 16S rRNA gene amplicons sequenced from whole fish samples after 24 hours of treatment (T0), 24 hours of recovery (T1), and 5 days of recovery (T5). A total of 2,161,882 quality-controlled, bacterial 16S rRNA gene amplicon sequences were analyzed from 45 larval zebrafish samples (Fig. S1A). Sequence variant analysis using QIIME2 and taxonomic classification resulted in the detection of 1,186 bacterial genera across 39 phyla, to sufficiently cover the estimated high diversity in the samples (Fig. S1B). Blank negative control samples were analyzed to confirm the absence of contamination relative to the zebrafish samples and a sequenced mock community yielded the expected sequencing proportions (Fig. S2). Proteobacteria was the dominant phylum in the larval zebrafish microbiota comprising 75 ± 17% of the samples, followed by Bacteroidota (9.6 ± 9.2%) and Firmicutes (5.0 ± 13%) (Fig. S3; values averaged across all samples). The largest group of 100 shared genera was common to all DMSO and loperamide samples, regardless of timepoint or treatment (Fig. S4B; black bar).

**Figure 1.**
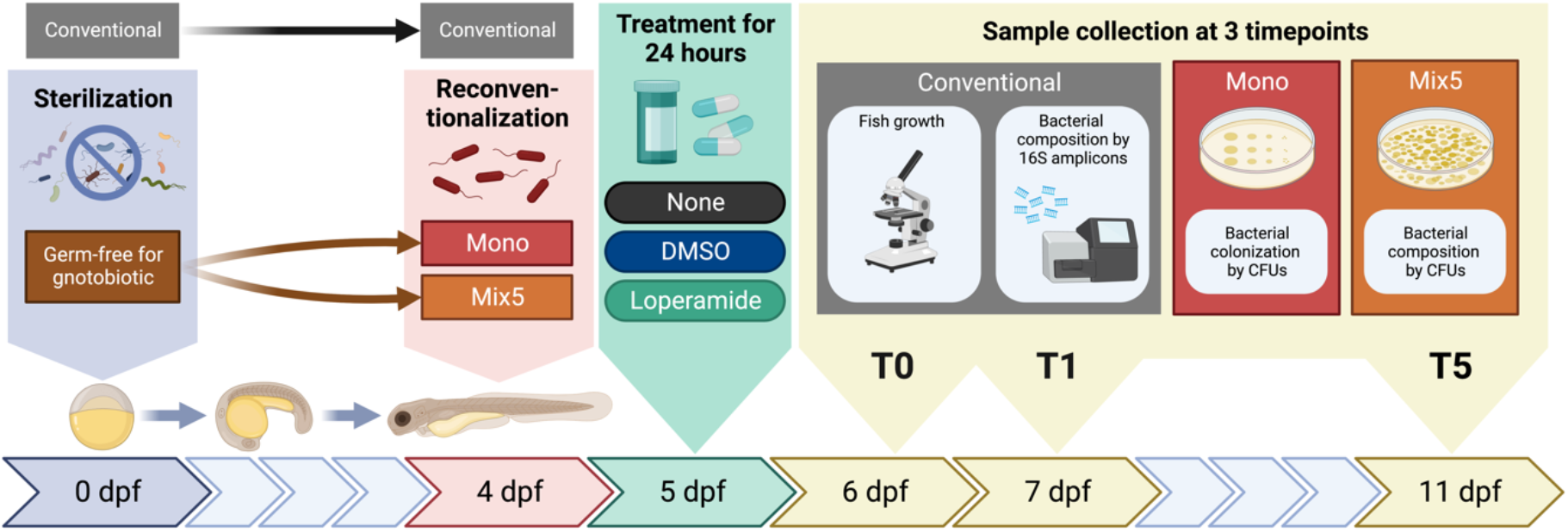
Experimental scheme of the larval zebrafish assays and sample collection. Conventional, mono-conventionalized, or Mix5-conventionalized larval zebrafish were exposed at 5 dpf to water (control), DMSO (control) or 10 mg/L loperamide hydrochloride (treated) for 24 hours, then transferred to water at 6 dpf. Samples were collected at 6 dpf, 7 dpf, and 11 dpf (T0, T1, T5) to measure fish growth and quantify bacterial community composition in all conditions (dpf = days post fertilization; CFUs = colony forming units).

Differences in the conventional zebrafish bacterial community composition were observed between the timepoints and treatment (Figs. 2, S3, S4, and S5). At T0 and T1, the beta-diversity of loperamide-treated fish microbiota was significantly different from the DMSO control (Figs. 2A, S5BC; adonis2 PERMANOVA R^2^ = 0.43, 0.51; p < 0.01). However, after 5 days of recovery (T5), the DMSO and loperamide-treated fish microbiota composition was not significantly different, indicating a recovery of microbiota composition once the treatment ended (Fig. 2A, S5D; adonis2 PERMANOVA R^2^ = 0.22; p > 0.05). This recoverable dysbiosis in microbiota composition induced by loperamide treatment was driven by a decrease in genus *Ensifer* and an increase in genus *Aeromonas* at T0 (Figs. 2B, S4A). At T1, there were major significant differences, affecting 37 different taxonomic groups, 6 of them with >1% abundance: a 4-log decrease in *Acidovorax* and significant enrichment of *Comamonadaceae, Acinetobacter, Flavobacterium, Oxalobacteraceae*, and *Rheinheimera* taxa (Fig. 2B). After 5 days of recovery at T5, there were no significantly different genera that were >1% abundant in the conventional zebrafish (Fig. 2B). Loperamide treatment also resulted in significantly decreased within-group beta-diversity compared to the DMSO control at T0 and T1, but not T5 (Fig. 2C). Despite the differences in bacterial composition and treatment regimen, growth and development of conventional zebrafish at all three timepoints was not affected by loperamide treatment (Fig. S6).

**Figure 2.**
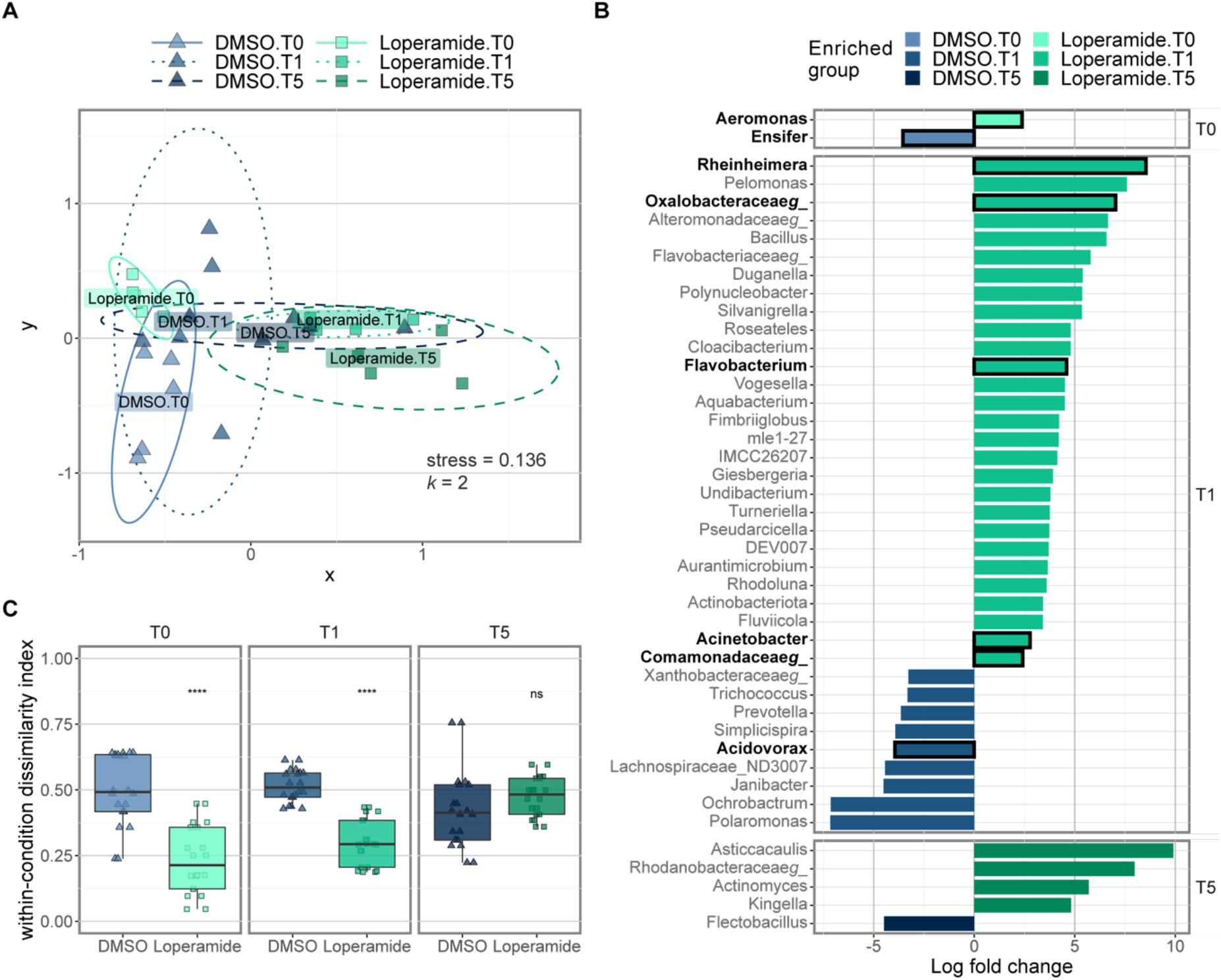
Loperamide affects conventional zebrafish microbiota as measured by 16S rRNA gene amplicons. **(A)** NMDS plot calculated using Bray-Curtis beta-diversity (k = 2) of percent normalized ASVs from 16S rRNA gene amplicons. Ellipse lines show the 95 % confidence interval (standard deviation). Stress = 0.136; adonis2 PERMANOVA R^2^ = 0.104; p = 0.029*. Only DMSO and loperamide-treated samples are shown. **(B)** Significant differentially abundant genera in loperamide-treated fish, compared to DMSO controls at each timepoint calculated using Limma (one-against-one) with conditions: relative log expression (RLE) normalized, effect log fold change >2, Benjamini & Hochberg adjusted p-value < 0.05 (n = 5 per condition). Genera that occur at mean percent abundance >1% are outlined in black and bold. **(C)** Beta-dispersion or within-condition dissimilarity index calculated using Bray-Curtis beta-diversity (n = 20). **** p<0.001 for Loperamide treatment, compared to DMSO. Wilcoxon test.

### Members of the conventional zebrafish community are inhibited or promoted by loperamide in vitro

Based on the changes observed in the conventional zebrafish bacterial community, 9 strains isolated from the zebrafish environment (conventional larvae or rearing water) and a *Flavobacterium* spp. were tested for their sensitivity to loperamide *in vitro* (Table S1). When grown in rich media in the presence of loperamide, the growth of 8/10 strains was significantly affected, while S2 *Variovorax gossypii* and S3 *Pseudomonas nitroreducens* were not affected (Fig. S7). One strain (S1 *Pseudomonas mosselii*) showed increased growth rate and carrying capacity in the presence of loperamide, compared to DMSO control. All other affected strains (7/10) showed no growth, delayed growth, slower growth rates or reduced carrying capacity when grown in media supplemented with loperamide (Fig. S7). In addition to growth, survival in water according to *in vivo* conditions was tested for the 10 strains by counting daily CFUs for three days of incubation. Survival of 6 out of 10 strains was not significantly affected by loperamide in these conditions: S1, S4, S5, S6, S9, and S10 (Fig. 3). Three strains (S2, S3, S8) showed increased survival in the presence of loperamide, while S7 *Aeromonas veronii* was the only strain with significantly inhibited survival at 24h.

**Figure 3.**
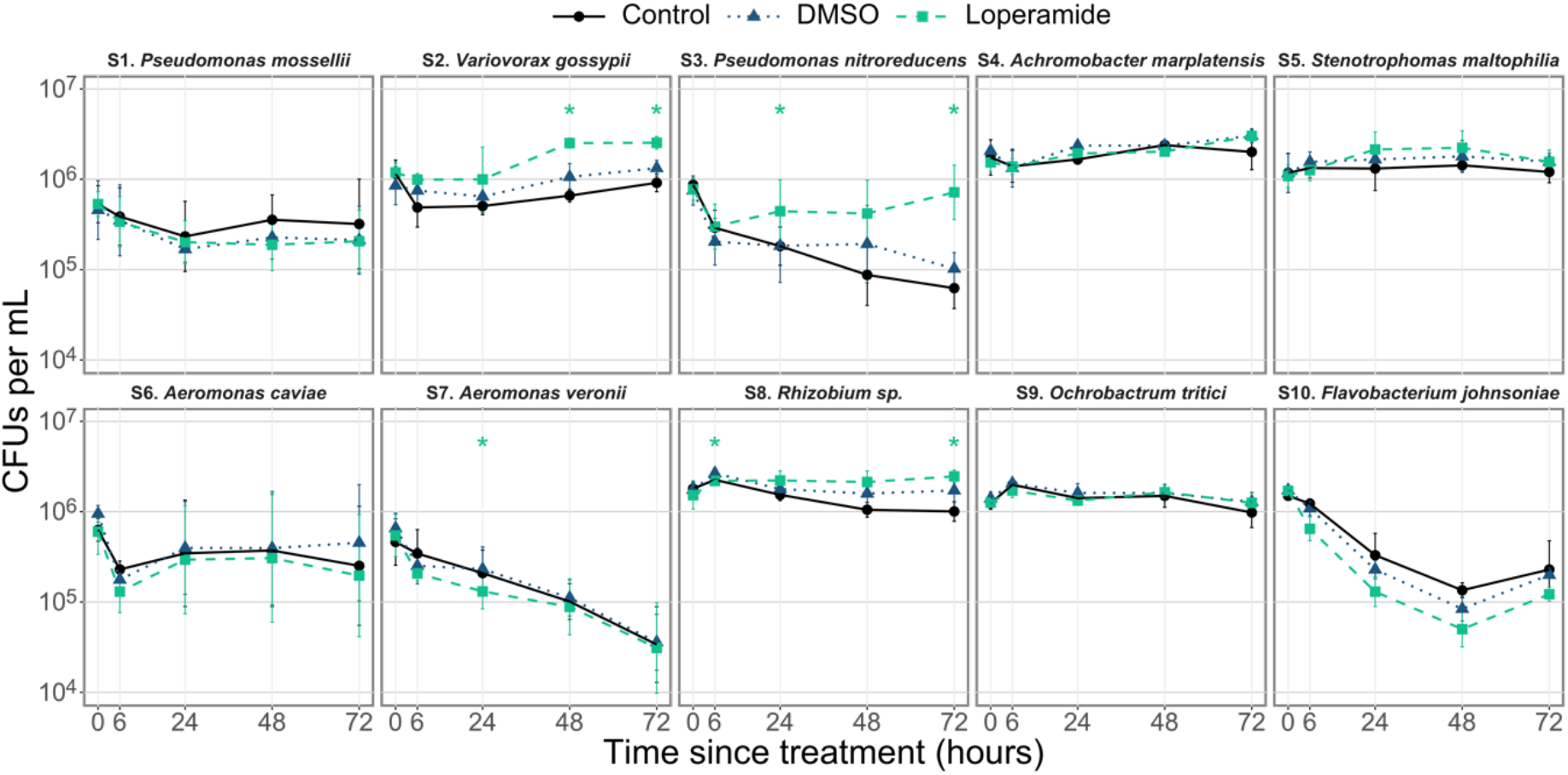
*In vitro* survival in water of zebrafish-associated bacterial strains is affected by loperamide. Survival in water for 72 hours after inoculation at 10^6^ CFUs/mL (mean ± standard deviation per condition is shown, n = 6-12: 2-4 independent assays of 3 biological replicates). * p<0.05 for loperamide treatment, compared to DMSO. Wilcoxon test. Note log scale on y-axis.

### Individual bacterial colonization of mono-reconventionalized larval zebrafish is strain-specific and affected by loperamide

In order to test the zebrafish colonization capacity of bacteria and the loperamide effects *in vivo*, 10 bacterial strains were individually added to reconventionalize GF fish and then sampled at T0, T1, and T5 for whole fish CFU counts. All bacterial strains colonized the zebrafish in control conditions at 6 dpf after 2 days of re-conventionalization at 10^3^ to 10^6^ CFUs per larvae (Fig. 4A). S4 *Achromobacter marplatensis* had the highest bacterial colonization capacity at a mean of 2.2 × 10^5^ CFU/fish, while the bacterial load of larvae reconventionalized with non-autochthonous S10 *Flavobacterium johnsoniae* was only 4.6 × 10^3^ CFU/fish. Overall bacterial colonization of the zebrafish was on average 10- to 100-fold lower than the number of CFUs per mL in the water at this time with colonization efficiencies of 0.7 – 52 % (Fig. S8A). Strain S7 *A. veronii* displayed the highest colonization efficiency with a mean of 1.1 × 10^5^ CFUs/mL in the water, compared to 5.9 × 10^4^ CFUs per fish (efficiency = 52.9 %). Conversely, strains S8, S5, and S2 had colonization efficiencies of ∼1% with ∼10^6^ CFUs/mL in the water, compared to ∼10^4^ CFUs per fish (Fig. S8A). Despite these large strain-specific differences in colonization efficiency, overall bacterial colonization per fish correlated with number of bacteria in the water at the time of sampling (Fig. S8B; R^2^ = 0.69, p = 0.03*).

**Figure 4.**
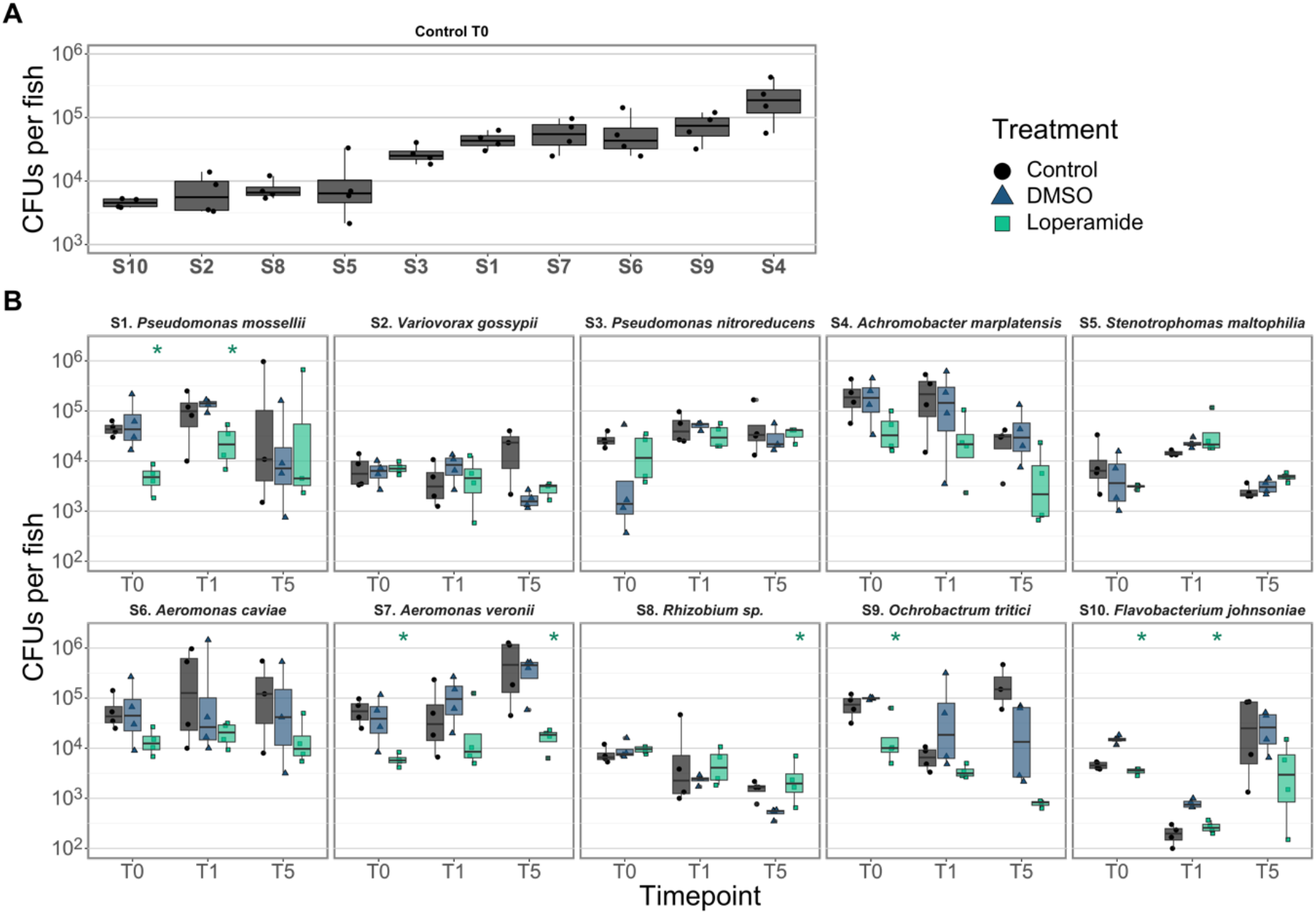
Loperamide can increase or reduce mono-reconventionalized zebrafish colonization. CFUs per fish of mono-reconventionalized fish in **(A)** control conditions at T0 (6 dpf) ordered by colonization capacity and **(B)** after exposure to loperamide at 3 timepoints (n = 4 fish). Each point represents a single zebrafish (mean of 3 technical replicates). * p<0.05 for loperamide treatment, compared to DMSO. Wilcoxon test. Note log scale for y-axis.

The addition of loperamide led to a reduction or increase in larval zebrafish bacterial load for half of the assayed strains. Five strains were not significantly affected by loperamide in mono-reconventionalized zebrafish: S2, S3, S4, S5, and S6. Colonization of larvae exposed to S1 *P. mosselii* or S10 *F. johnsoniae* was significantly reduced in the presence of loperamide at T0 and T1, but recovered to match DMSO-level colonization by T5 (Fig. 4B; p < 0.05). S7 *A. veronii* and S9 *Ochrobactrum tritici* bacterial load was reduced at all timepoints with loperamide treatment. One strain (S8 *Rhizobium* sp.) showed higher colonization only at T5 after loperamide treatment (Fig. 4B; p < 0.05). These strain-specific colonization changes due to loperamide confirm inhibition or promotion of bacteria *in vivo*, in addition to the host-exclusive effects of the molecule. A summary of how loperamide affects *in vitro* growth and survival, and *in vivo* mono-colonization of all strains is detailed in Table 1.

**Table 1.** Summary of *in vitro* effects of loperamide on zebrafish strains. Significant changes in growth in media, survival in water, and *in vivo* zebrafish colonization for loperamide-treated, compared to DMSO control.

| Strain | Growth in media | Survival in water | Zebrafish mono colonization |
| --- | --- | --- | --- |
| S1. *Pseudomonas mossellii* | Promoted | No effect | Reduced at T0 and T1, then recovery |
| S2. *Variovorax gossypii* | No effect | Increased from 6 hours | No effect |
| S3. *Pseudomonas nitroreducens* | No effect | Increased from 24 hours | No effect |
| S4. *Achromobacter marplatensis* | Reduced | No effect | No effect |
| S5. *Stenotrophomas maltophilia* | Reduced | No effect | No effect |
| S6. *Aeromonas caviae* | Inhibited | No effect | No effect |
| S7. *Aeromonas veronii* | Inhibited | Decreased at 24 hours | Reduced at T0 and T5 |
| **S8. *Rhizobium* sp.** | Inhibited | Increased from 24 hours | Increased at T5 |
| **S9. *Ochrobactrum tritici*** | Inhibited | No effect | Reduced at T0 |
| **S10. *Flavobacterium johnsoniae*** | Inhibited | No effect | Reduced at T0 and T1, then recovery |

### Loperamide treatment induces expected dysbiosis in mix-reconventionalized gnotobiotic larval zebrafish

In order to evaluate how loperamide affects a multi-species bacterial community *in vivo*, germ-free zebrafish were reconventionalized with an equal mix of strains S1, S3, S5, S6, and S7. These strains were selected according to their varying sensitivities to loperamide *in vitro* and *in vivo*. Loperamide treatment did not significantly impact the total number of CFUs per mix-reconventionalized fish (Fig. 5A). However, the addition of loperamide induced an increase in S7 *A. veronii* and a decrease in S6 *A. caviae* at T0, relative to the DMSO control (Fig. 5B). Meanwhile, S3, S5, and S6 increased in loperamide-treated samples at T1. Finally, at T5 after 5 days of recovery, S5 *S maltophila* and S7 *A. veronii* were the most abundant strains. (Fig. 5B). These changes in the proportion of each strain per fish reflect *in vitro* sensitivity to loperamide and changes measured in conventional fish during loperamide treatment.

**Figure 5.**
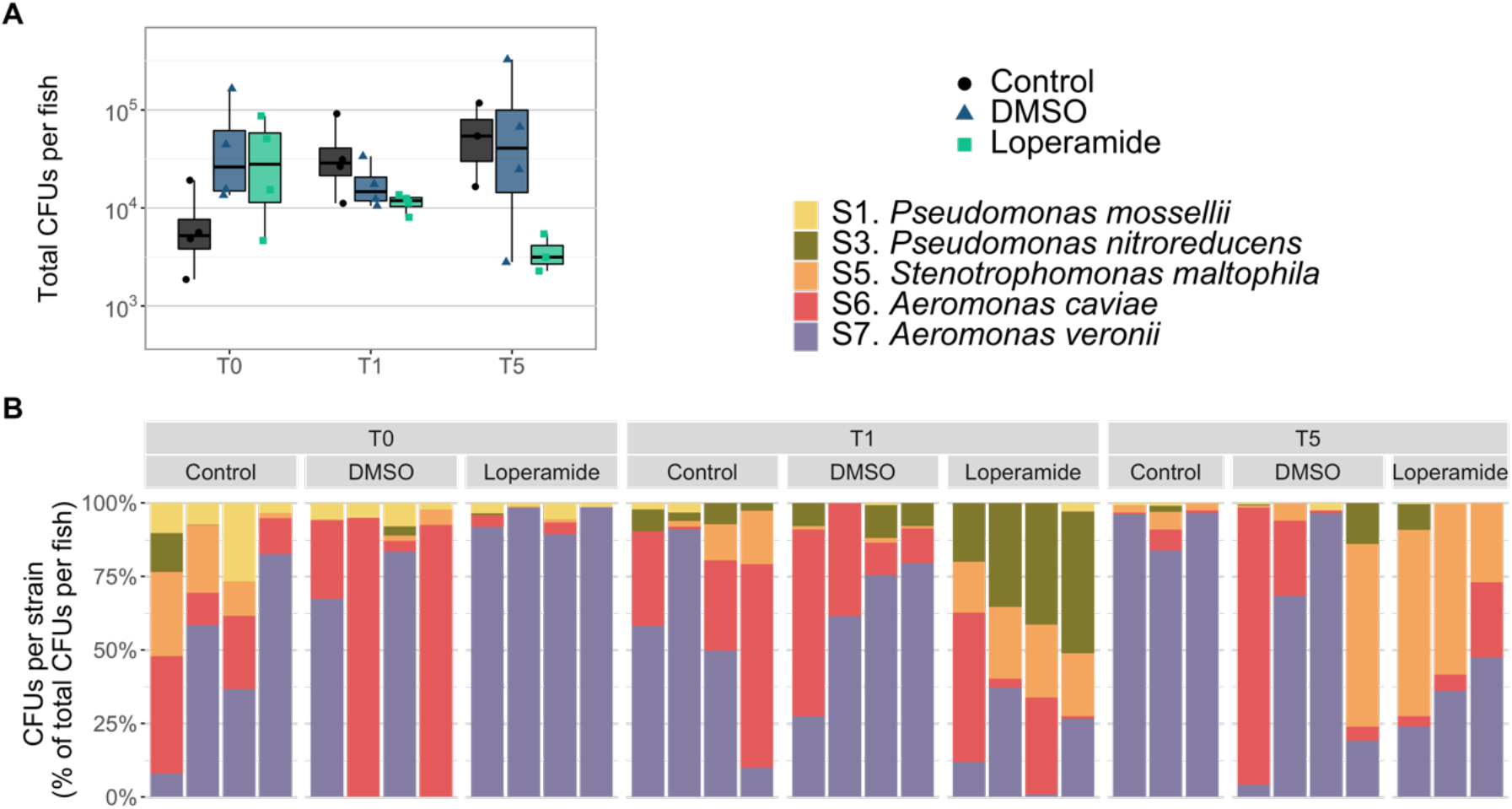
Loperamide affects mix-reconventionalized gnotobiotic zebrafish bacterial load and composition. **(A)** Total CFUs per fish of mix-reconventionalized fish after exposure to loperamide at 3 timepoints (n = 3-4 fish). Each point represents a single zebrafish (mean of 3 technical replicates). No significant changes were found for loperamide treatment, compared to DMSO. Wilcoxon test. Note log scale for CFUs. **(B)** Percent abundance of each strain per mix-reconventionalized fish. Each bar is an individual fish sample.

Differences in strain-specific colonization efficiency in zebrafish individually reconventionalized with these 5 strains may have contributed to loperamide-independent effects on the mix-reconventionalized bacterial colonization (Fig. S8). We compared the mix-reconventionalized bacterial composition with the sum of mono-reconventionalized bacterial abundances for S1, S3, S5, S6, and S7 (Fig. S9). This comparison of the mono means to the mix showed that the composition of the mix-reconventionalized fish was different from the sum of the mono-conventionalized fish in all conditions (Fig. S9AB). Therefore, inter-bacterial competition in the mix-reconventionalized fish also contributed to changes in community composition, in addition to host selection and bacterial inhibition by loperamide. Comparison of the CFUs per strain in mono-reconventionalized fish to mix-reconventionalized fish also showed increased colonization for each strain in mono-than when part of a mix, regardless of timepoint or treatment and despite the increased number of bacteria added (Fig. S9CD). Even in control conditions, each strain colonized 10-10000 times higher when added alone than when added as part of a mix (Fig. S9CD).

Further comparison of the bacterial composition in conventional and gnotobiotic zebrafish focused on changes in alpha-diversity after loperamide treatment and during recovery. Loperamide-treated conventional fish alpha-diversity measured by Simpson’s Index significantly decreased after 24 hours of loperamide treatment (T0; p<0.05), then increased after 24 hours of recovery (T1) and stayed similar to control diversity at T5 days post-treatment (Fig. 6A). This decrease in diversity was confirmed by the lower number of ASVs (richness) detected in the loperamide-treated samples at T0 (Fig. S1B, first panel). Similarly, the alpha-diversity of loperamide-treated mix-reconventionalized gnotobiotic zebrafish decreased at T0, significantly increased at T1 (p<0.05), and recovered to match the control at T5 (Fig. 6B). These results show that in both natural and synthetic zebrafish bacterial communities, loperamide induced a significant, but recoverable, dysbiosis and associated loss in diversity.

**Figure 6.**
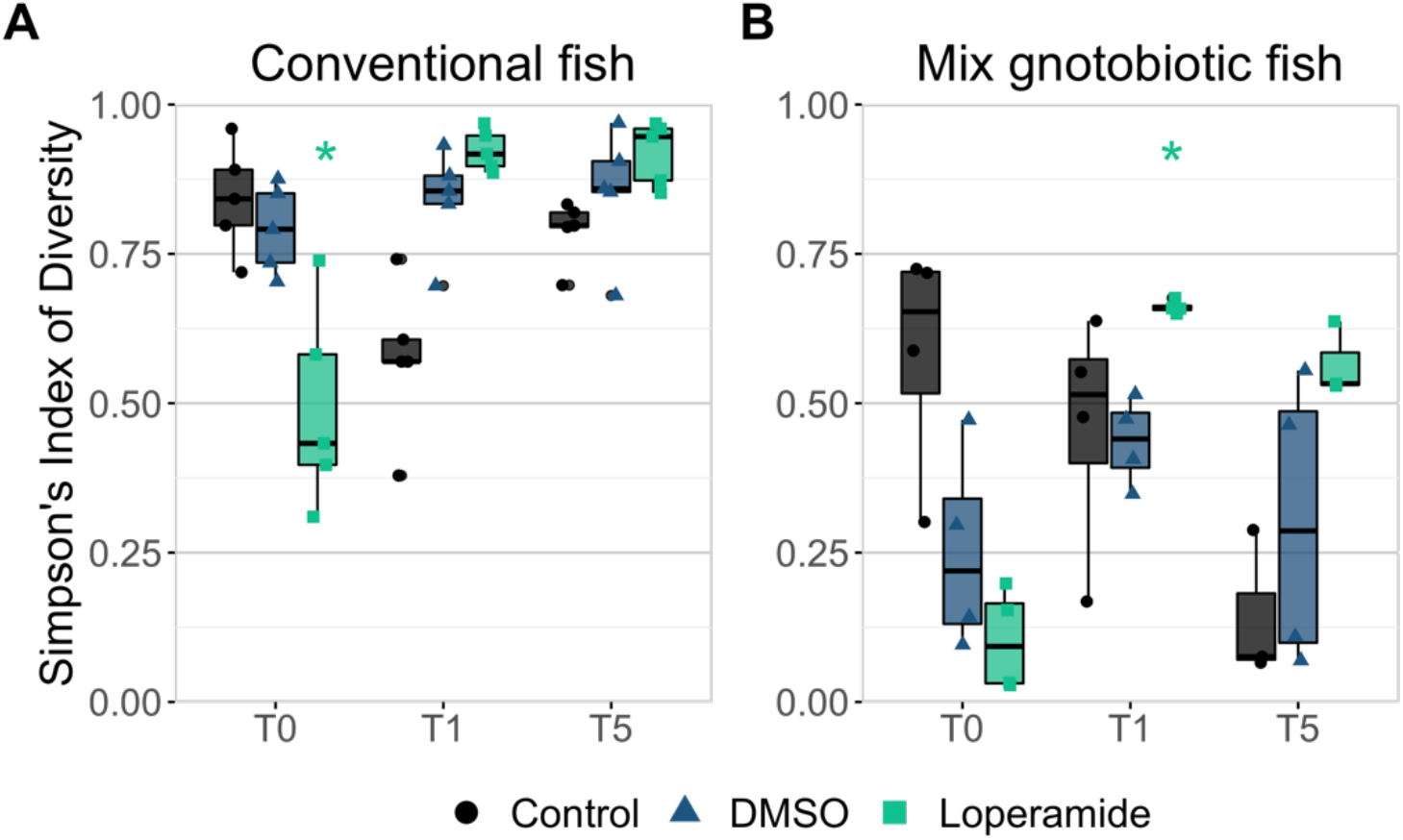
Alpha-diversity of conventional and gnotobiotic zebrafish decreases but recovers after loperamide treatment. Simpson’s index of diversity calculated at each timepoint for control water, DMSO, and loperamide-treated samples for **(A)** 16S rRNA gene amplicon data at the ASV level in conventional fish (n = 5 fish) and **(B)** CFUs per strain in mix-reconventionalized fish (n = 3-4 fish). Each point represents a single zebrafish with boxplots shown per condition. * p<0.05 for loperamide treatment, compared to DMSO. Wilcoxon test.

## Discussion

Understanding the impact of non-antibiotic drugs on host-associated microbiota is critical for sustaining health in humans as well as animal models. In this study, we evaluated the effects of loperamide, a widely prescribed anti-diarrheal compound also used as a tool to study the impact of bowel dysfunctions in animal models. Using conventional and gnotobiotic zebrafish, we showed that loperamide directly induced significant but recoverable dysbiosis by broad-range inhibition. The effects of loperamide on zebrafish-associated bacteria characterized by growth, survival, and colonization capacity were strain-specific and changed in the presence of other bacteria or the zebrafish host.

Loperamide-induced decreases in microbiota alpha-diversity and beta-dispersion immediately after loperamide treatment. These changes were not permanent and initial alpha-diversity recovered within 24 hours after loperamide exposition, and within 5 days for beta-diversity. These results were consistent with a previous study in mice, in which loperamide was used to increase gastrointestinal transit time, but also led to alterations in the gut microbial community that were reversible after treatment interruption [58]. This dysbiosis was presumed to result from a reduction of peristaltic movement, but our results suggest that it could also be explained by the loperamide bactericidal activity [27, 28].

We found that the effects of loperamide treatment on zebrafish microbiota composition depended on a strain’s survival in water and colonization capacity. In conventional zebrafish, loperamide induced a significant increase in the *Aeromonas* genus at T0, but not at T1 or T5. In mono-reconventionalized fish, S6 *A. caviae* was not affected by loperamide, but S7 *A. veronii* showed impaired colonization despite its high colonization efficiency. S7 *A. veronii* was the only strain with inhibited growth and decreased survival in water over time, which may have contributed to its inability to recover colonization capacity after loperamide treatment. In the mix-reconventionalized fish, S7 *A. veronii* was the most abundant strain in the loperamide-treated zebrafish at T0, but significantly decreased at T1 and T5, consistent with its colonization in the mono-reconventionalized larvae and the conventional zebrafish composition. Other bacteria- or host-related factors induced by the presence of loperamide could explain reduced S7 *A. veronii* abundance, such as reduced feeding, chemokinesis, or motility [22, 59, 60]. Previous studies of gnotobiotic zebrafish colonization have demonstrated the strain-specific importance of chemotaxis and host gut motility for intestinal colonization [61, 62], bacterial motility and host cues with *A. veronii*. [60], and general induction of host immune responses or locomotive behavior [63–65]. In a mix-reconventionalized community, bacteria-bacteria interactions also contribute to changes in relative abundance, regardless of host factors. For example, the ecological niche left by S7 *A. veronii* due to direct inhibition or decreased intestinal peristalsis from loperamide treatment, could explain why S3 *P. nitroreducens* showed a significant increase at T1 only in loperamide-treated samples.

Interestingly, loperamide did not increase bacterial load at the measured timepoints in our study. Similar results were also obtained in loperamide-induced constipation model in rats [66]. This may be due to colonization constraints imposed by loperamide toxicity, the larval fish size, or nutrient limitations, since previous studies of gnotobiotic zebrafish have also not detected more than 10^6^ CFUs/larvae [22, 63, 64]. Our study is limited to bulk culturable CFUs per fish associated with 10 bacterial strains at 3 timepoints. Future studies should investigate the localization and quantification of transit time of fluorescently tagged bacteria to further understand intestinal-specific changes upon loperamide treatment.

In all previous studies where loperamide-induced constipation has been considered to affect the host microbiome, these changes have been attributed to decreased stool frequency and increased colonic contractions by inhibition of intestinal water secretion and colonic peristalsis, which extends the fecal evacuation time and delays the intestinal luminal transit rate [15, 67]. However, our results demonstrated that the changes in microbiota composition and diversity are also partially due to strain-specific bacterial inhibition or promotion by the loperamide exposure. In addition to the zebrafish-associated strains studied here, loperamide exhibits bactericidal activity against diverse host-associated microbes including mycobacterial strains (e.g. *Mycobacterium tuberculosis*) and *Staphylococcus aureus*, but not *Escherichia coli* [27, 68]. These microbes are members of the human and vertebrate microbiome that may be directly affected by loperamide treatment, resulting in unforeseen microbiota modulation [69].

Prior studies of loperamide-induced gastro-intestinal disorders determined that various treatments restore host health and improve the associated symptoms (i.e. constipation or gut transit time). For example, konjac oligo-glucomannan alleviates defecation infrequency and suppressed the growth of *Bacteroides* in mice [70], raffino-oligosaccharide improved gastro-intestinal transit rate and reduced the serum levels of vasoactive intestinal peptide in mice [71], and probiotics improved constipation by altering metabolite, amino acid, inflammatory cytokines, and/or neurotransmitter abundances in rats [20, 72, 73]. In all of these studies, the effect of treatment and changes in host physiology were inferred to constipation or the relevant model phenotype. However, all of these described effects may also be attributed to ancillary microbiota modulations. The perturbation of host microbiomes is frequently described to cause significant changes in host metabolite and peptide abundances, immune response, and physiology and health [1, 74, 75]. Our results indicate that animal models using loperamide to study bowel dysfunction and constipation cannot distinguish the effects of loperamide on host function from the effects of microbiota modulation by loperamide.

## Conclusions

In summary, our results demonstrate that loperamide induces significant changes in the microbiota, which may influence experimental outcomes especially if the host immune system or behavior are considered. As a common medication used to alleviate diarrhea and bowel disorders in humans, loperamide is also likely to produce under-studied antibiotic effects on intestinal microbiota. This emphasizes the need to better characterize relationships between host physiological changes, microbial community structure, and disease or dysbiosis states.

## Acknowledgements

We are grateful to Jean-Pierre Levraud, Yuliaxis Ramayo and Erin Witkop for critical reading of the manuscript, and Bianca Audrain for assistance with obtaining the animal ethical authorization. This work was supported by grants from the French Government’s Investissement d’Avenir program, Laboratoire d’Excellence “Integrative Biology of Emerging Infectious Diseases” (grant n°ANR-10-LABX-62-IBEID) and by the Fondation pour la Recherche Médicale (grant DEQ20180339185). RJS was supported by a grant from the Philippe Foundation. Sequencing was performed by G M. Haustant, L. Lemée, Biomics Platform, C2RT, Institut Pasteur, Paris, France, supported by France Génomique (ANR-10-INBS-09-09) and IBISA. The graphical abstract and Figure 1 were created with BioRender.com.

## Authors’ Contributions

RJS, JMG, and DPP contributed conception and design of the study. RJS and DPP performed the experiments. SB and ND performed the zebrafish imaging assays. RJS analyzed the data and wrote the first draft of the manuscript. All authors contributed to manuscript revision, read and approved the submitted version.

## Competing Interests

The authors declare no competing interests.

## Ethics Statement

All animal experiments described in the present study were conducted at the Institut Pasteur according to European Union guidelines for handling of laboratory animals (http://ec.europa.eu/environment/chemicals/lab_animals/home_en.htm) and authorized by the Institut Pasteur Animal Health and Care Committees under permit #dap220109.

## Availability of Data and Material

The raw 16S rRNA gene amplicon sequences generated for this study can be found in the NCBI Sequencing Read Archive in BioProject no. PRJNA908751. All other raw data, processed sequencing files, QIIME2 artifacts, and scripts to reproduce the figures in the manuscript are available in the Zenodo repository, https://doi.org/10.5281/zenodo.7415697 [48].

## Notes

### Competing Interest Statement

The authors have declared no competing interest.

