## Supplementary Table S1 Supplementary Figures S1 to S9 for "Anti-diarrheal drug loperamide induces dysbiosis in zebrafish microbiota via bacterial inhibition"

### Supplementary materials for:
#### *Anti-diarrheal drug loperamide induces dysbiosis in zebrafish microbiota via bacterial inhibition*

Rebecca J. Stevick, Sébastien Bedu, Nicolas Dray, Jean-Marc Ghigo and David Pérez-Pascual.

Institut Pasteur Université Paris Cité, CNRS UMR 6047, Genetics of Biofilms Laboratory, Paris F-75015, France.

This PDF file includes:

Supplementary Table S1

Supplementary Figures S1 to S9

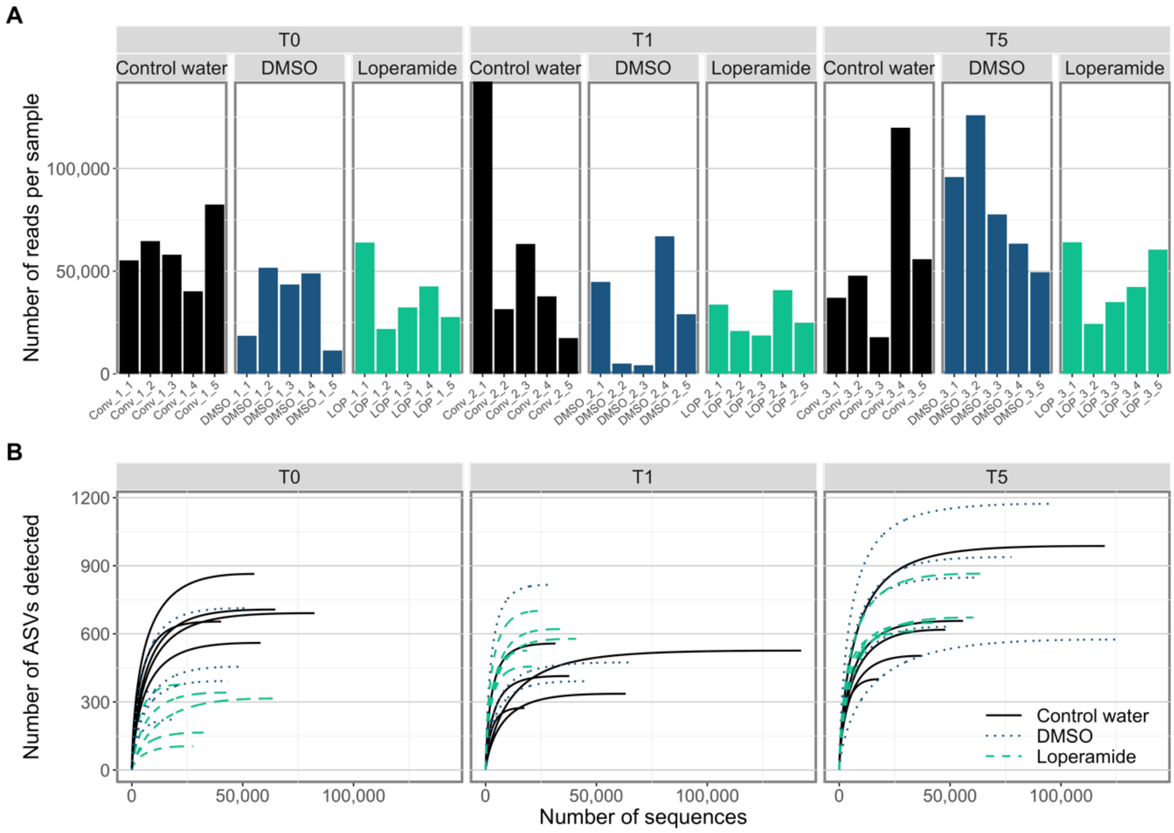


**Figure S1. Sequencing depth and coverage for 16S rRNA gene amplicon data from conventional zebrafish larvae. (A)** Number of quality-controlled bacterial sequences per sample. **(B)**. ASV rarefaction curves of all zebrafish larvae samples separated by timepoint and colored by treatment group (n = 5 fish per condition).

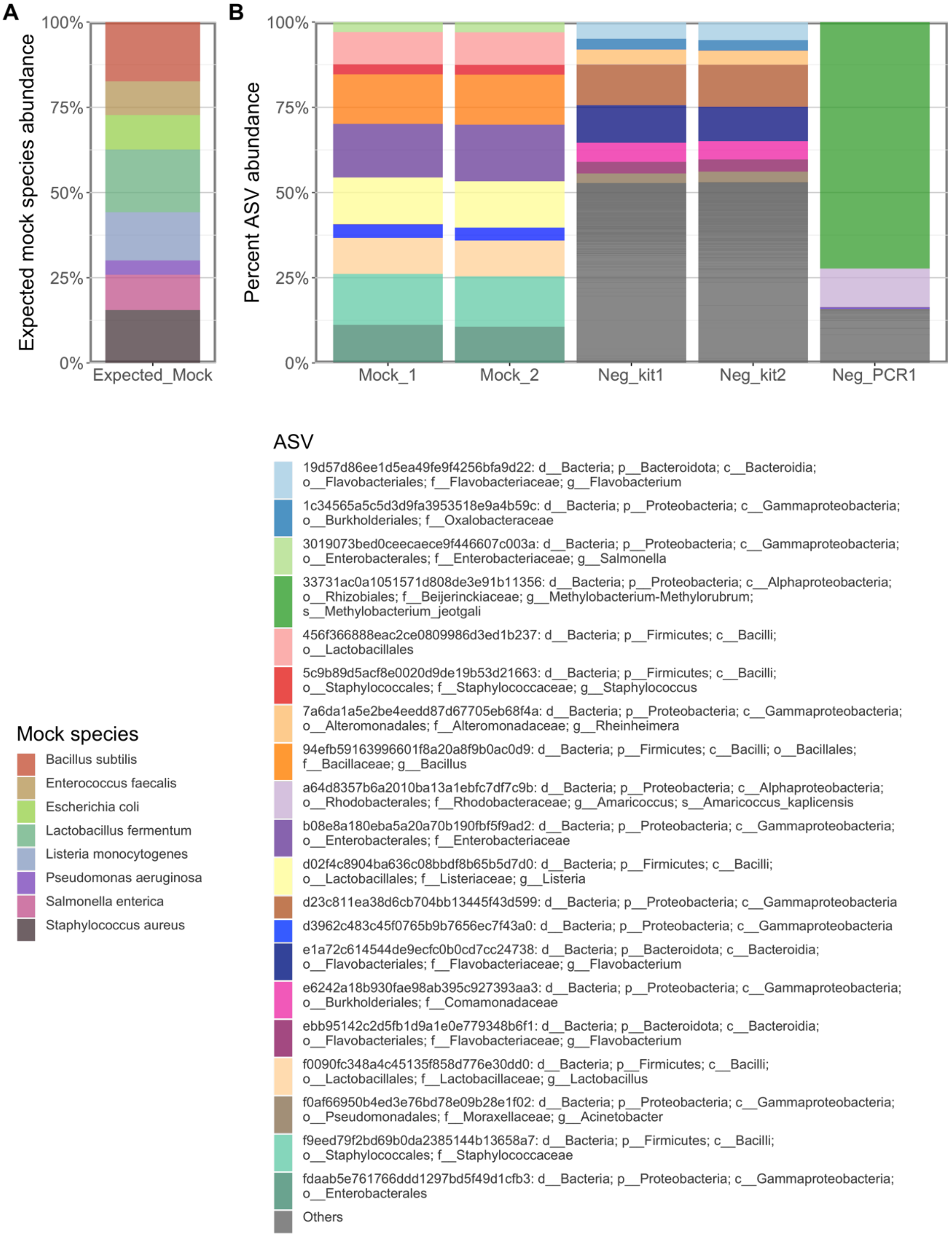


**Figure S2. Controls for 16S rRNA amplicon sequencing data: blanks and mock community. (A)** Expected mock community composition based on 16S rRNA gene copy number and relative percent abundance of each strain. **(B)** Relative percent abundance of ASVs for each negative or positive sequencing control sample: Zymo mock community standard D6305 (Mock_1 and Mock_2), negative DNA extraction (Neg_kit1 and Neg_kit2), negative PCR amplification (Neg_PCR1). The top 20 most abundant ASVs are shown, with all others grouped in the grey "Others" category.

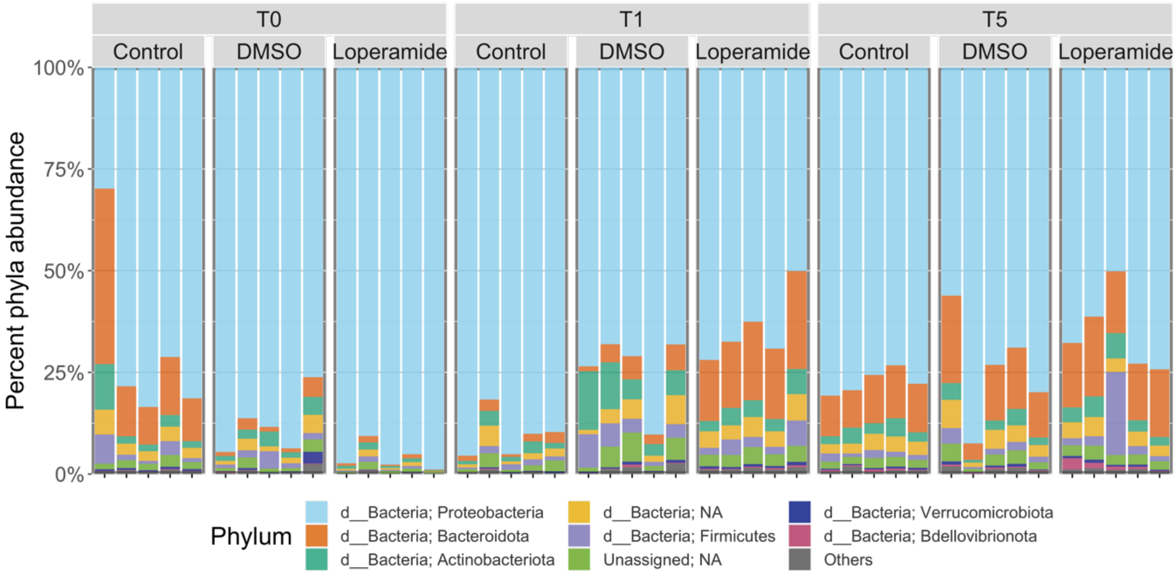


**Figure S3. 16S rRNA gene amplicon relative abundances at the phylum level. (A)** Bar plot of percent phylum abundance per sample. The top 8 most abundant phyla are shown with the others grouped into "Others" (n = 5 fish per condition).

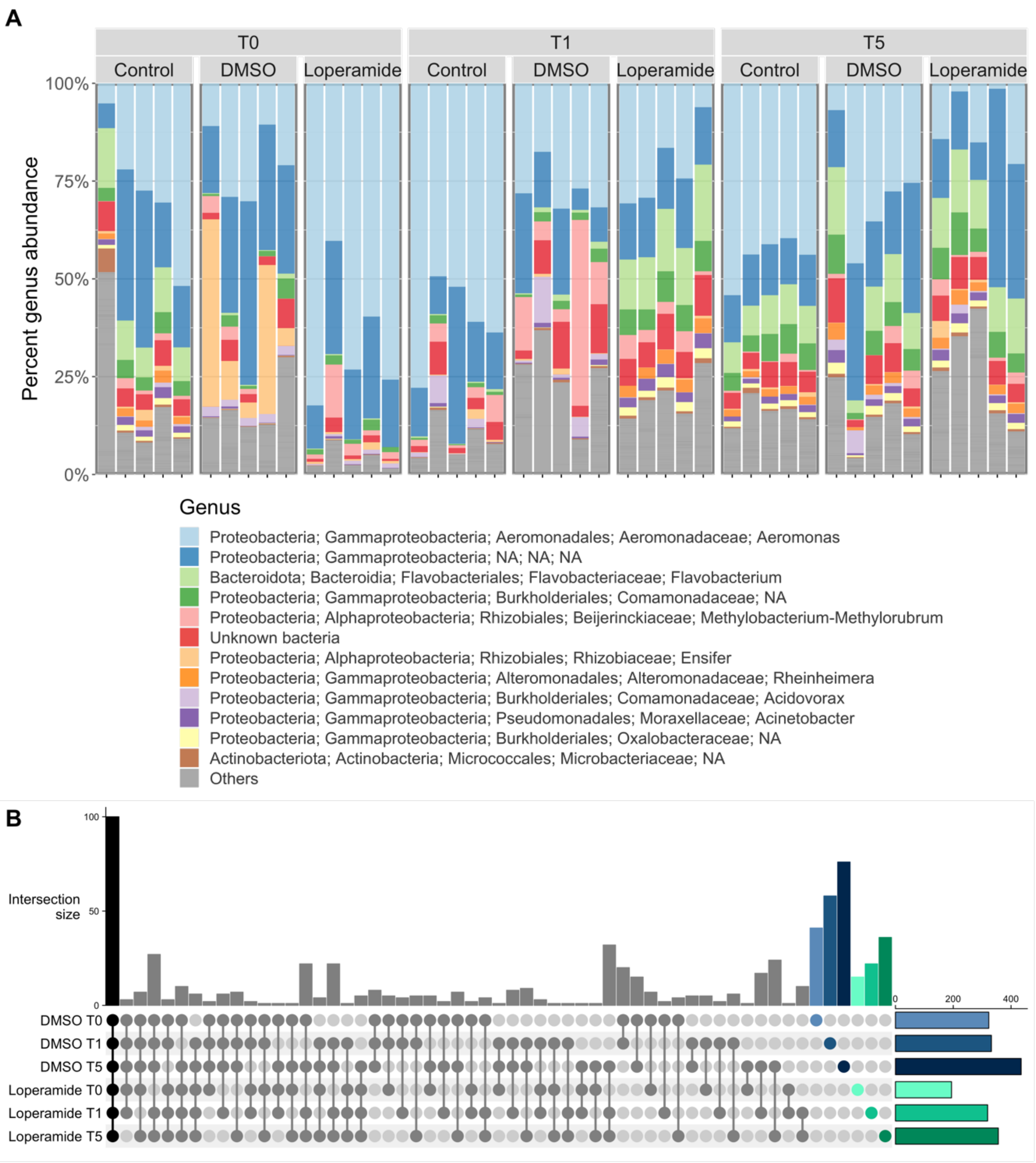


**Figure S4. 16S rRNA gene amplicon relative abundances at the genus level. (A)** Bar plot of percent genus abundance per sample. The top 12 most abundant genera are shown with the others grouped into "Others" (n = 5 fish per condition). **(B)** Number of bacterial genera shared between DMSO and Loperamide-treated samples at each timepoint (vertical bars). The total number of genera detected in each group is shown in the horizontal bar plot on the right.

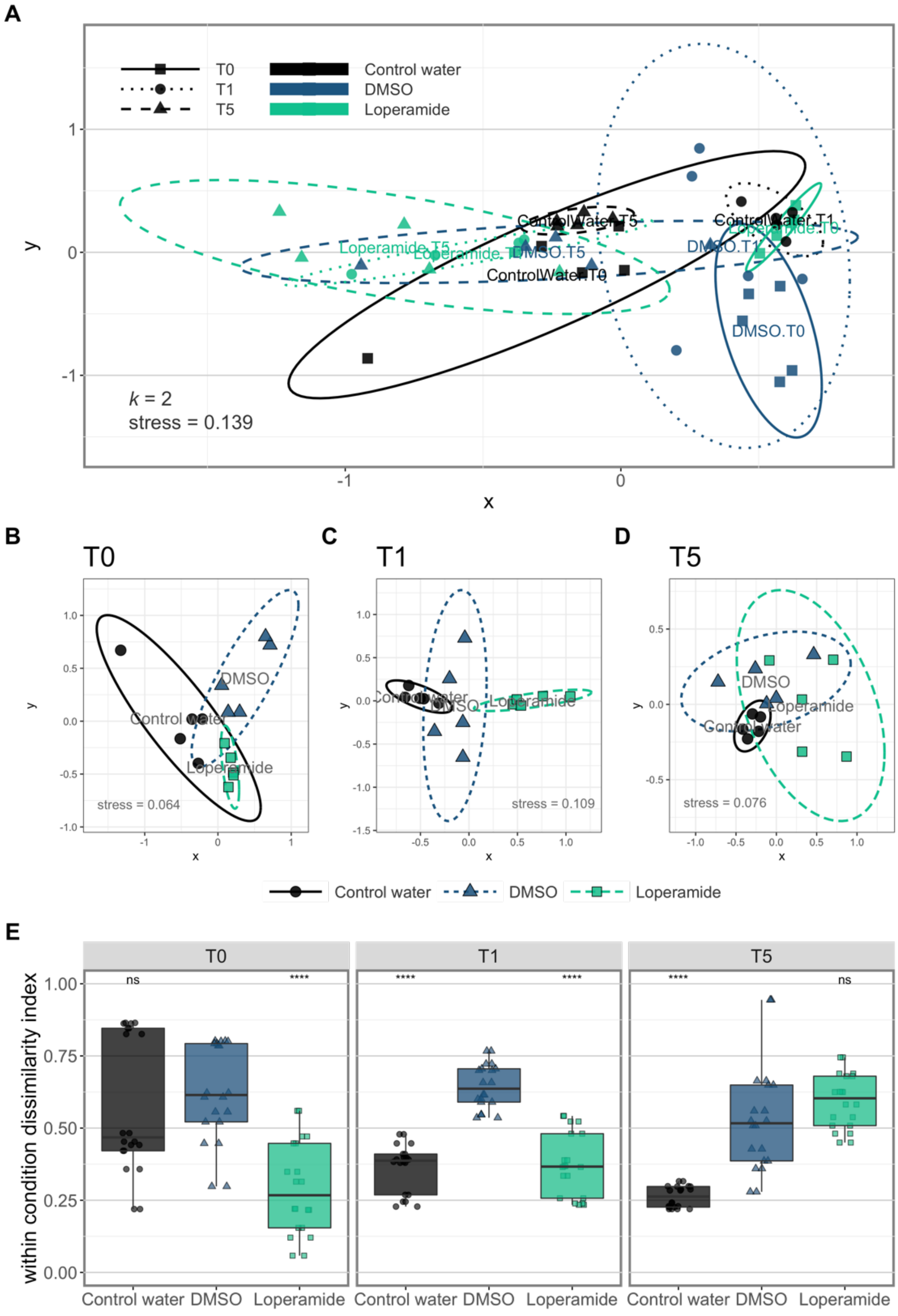


**Figure S5. Beta-diversity metrics of 16S rRNA gene amplicons sequenced from conventional zebrafish. (A)** NMDS plot calculated using Bray-Curtis beta-diversity (k=2) of percent normalized ASVs from 16S rRNA gene amplicons for all water control, DMSO control, and loperamide-treated samples. Ellipse lines show the 95 % confidence interval (standard deviation). Stress = 0.139 (n = 5 fish per condition). **(B-D)** NMDS plot calculated using Bray-Curtis beta-diversity (k=2) of percent normalized ASVs from 16S rRNA gene amplicons at each timepoint **(B)** T0 6 dpf, **(C)** T1 7 dpf, and **(D)** T5 11 dpf. The stress in indicated on each plot. **(E)** Beta-dispersion or within-condition dissimilarity index calculated using Bray-Curtis beta-diversity (n =15; 3 treatment groups for each of 5 samples per condition). **** $p<0.001$ for Loperamide treatment, compared to DMSO. Wilcoxon test.

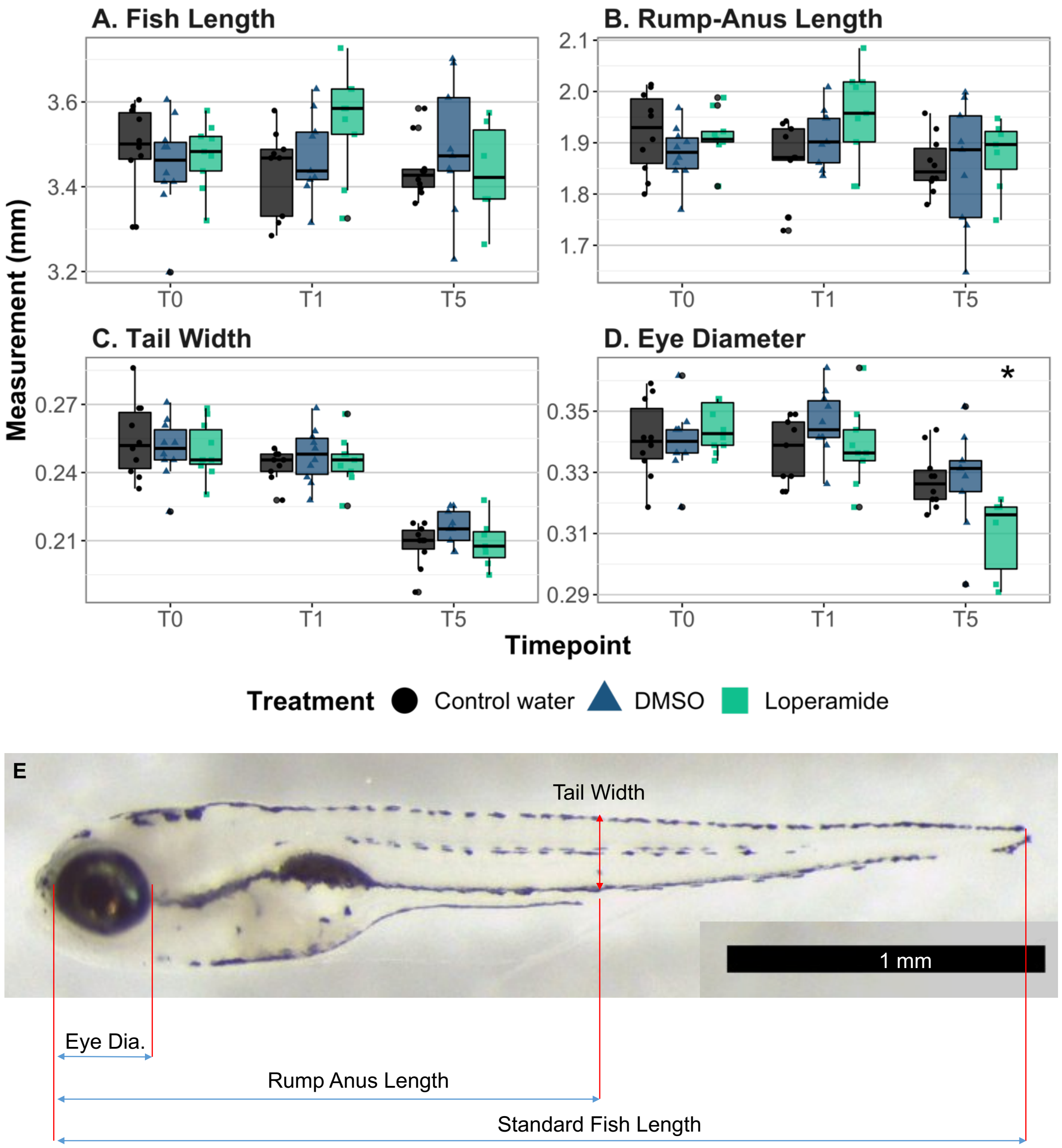


**Figure S6. Growth parameters measured for zebrafish larvae. (A)** Fish length, **(B)** rump-anus length, **(C)** tail width, and **(D)** eye diameter measurements of larval zebrafish at 6 dpf, 7 dpf, and 11 dpf (T0, T1, T5 after 24-hour treatment). All measurements are shown in millimeters (n = 10 fish per condition). * $p<0.05$ for Loperamide treatment, compared to DMSO. Wilcoxon test. The only significant difference is in (D) Eye diameter at T5. **(E)** Example fish image with the four measurements indicated and scale bar.

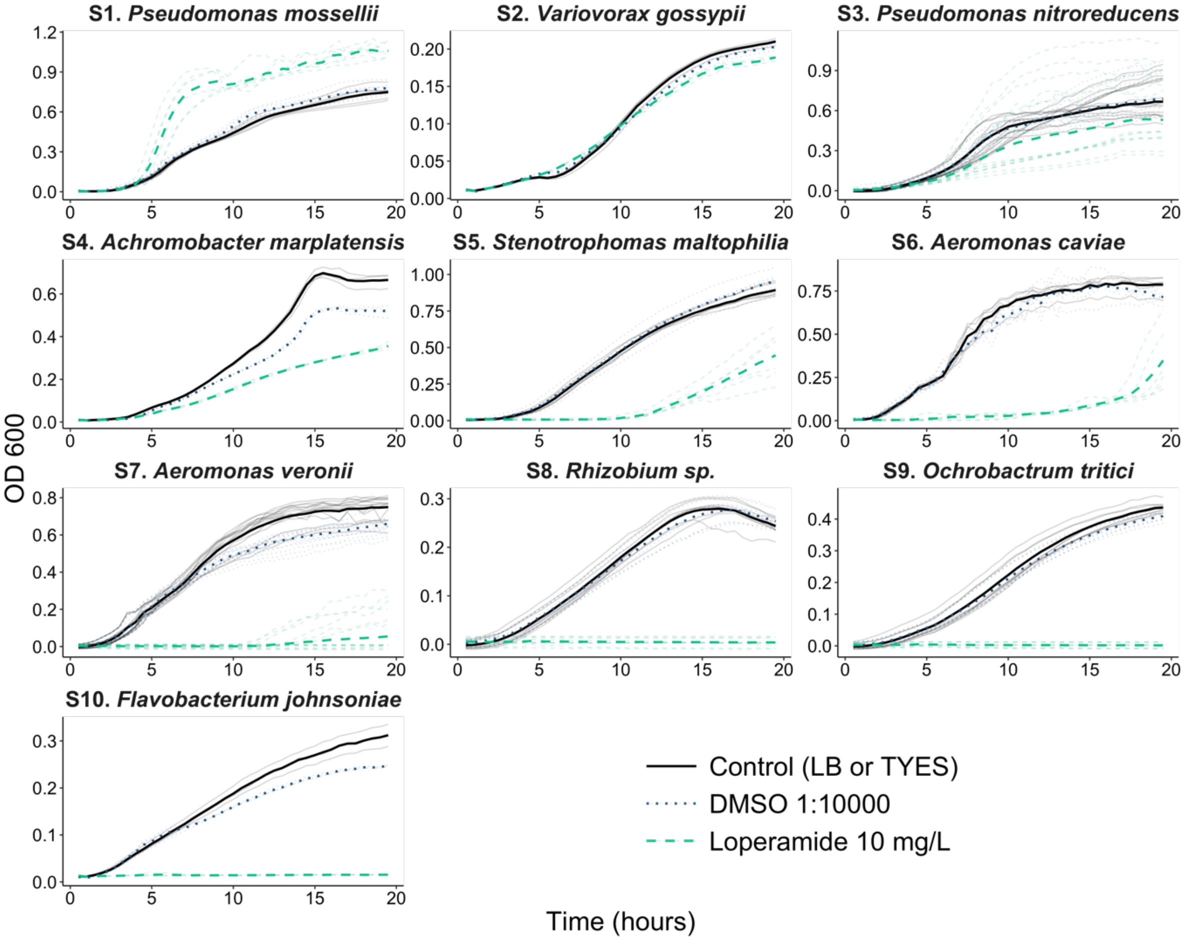


**Figure S7. Growth curves of zebrafish-associated bacterial strains exposed to loperamide.** Growth curves of 10 strains isolated from the fish environment or environmental *Flavobacterium* spp. (additional strain details are in Table S1). The thick line represents the mean of all biological replicates (n=3-8). Each thin line represents a biological replicate (mean of 3 technical replicates). Every condition was repeated at least twice.

23

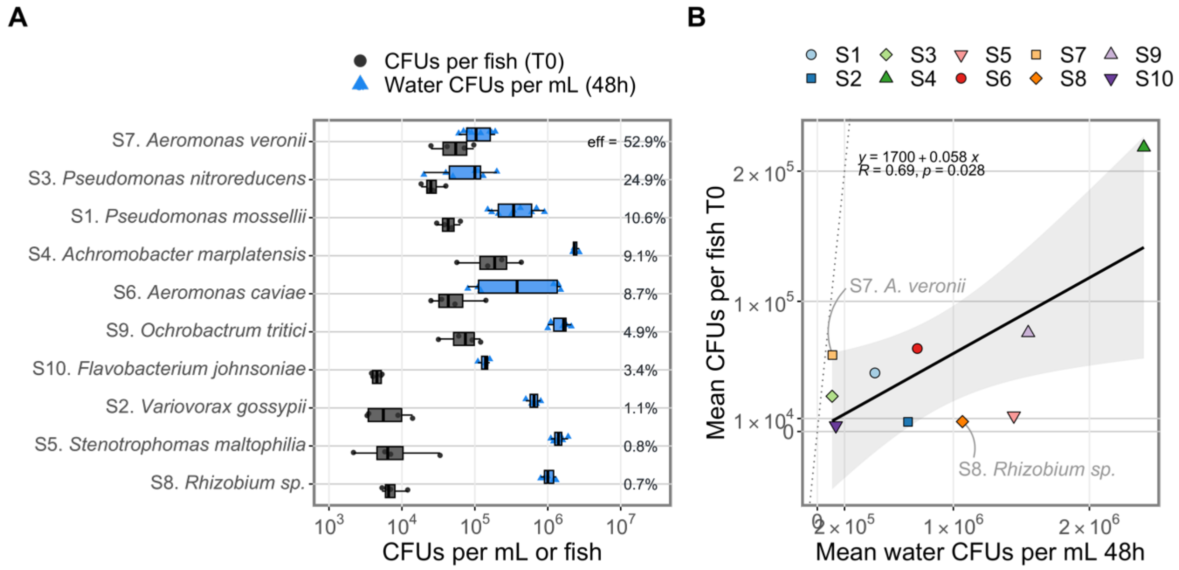


**Figure S8. Comparison of bacterial load in water and zebrafish colonization capacity in control conditions. (A)** Boxplots showing CFUs per mL in water at 48 h and CFUs per fish at T0 (6 dpf) for each of the 10 bacterial strains ordered by colonization efficiency. The value indicated on the plot is the colonization efficiency, calculated by *Colonization efficiency = CFUs per Fish / Water CFUs per mL * 100.* Note log scale on x-axis. **(B)** Correlation between bacterial load in water with zebrafish colonization efficiency. The grey dotted line indicates the 1:1 line. The regression line is indicated by the solid black line and the fitted equation, $R^2$ and p-value are shown in the top left corner. The strains with the lowest (S8) and the highest (S7) colonization efficiency are highlighted.

24

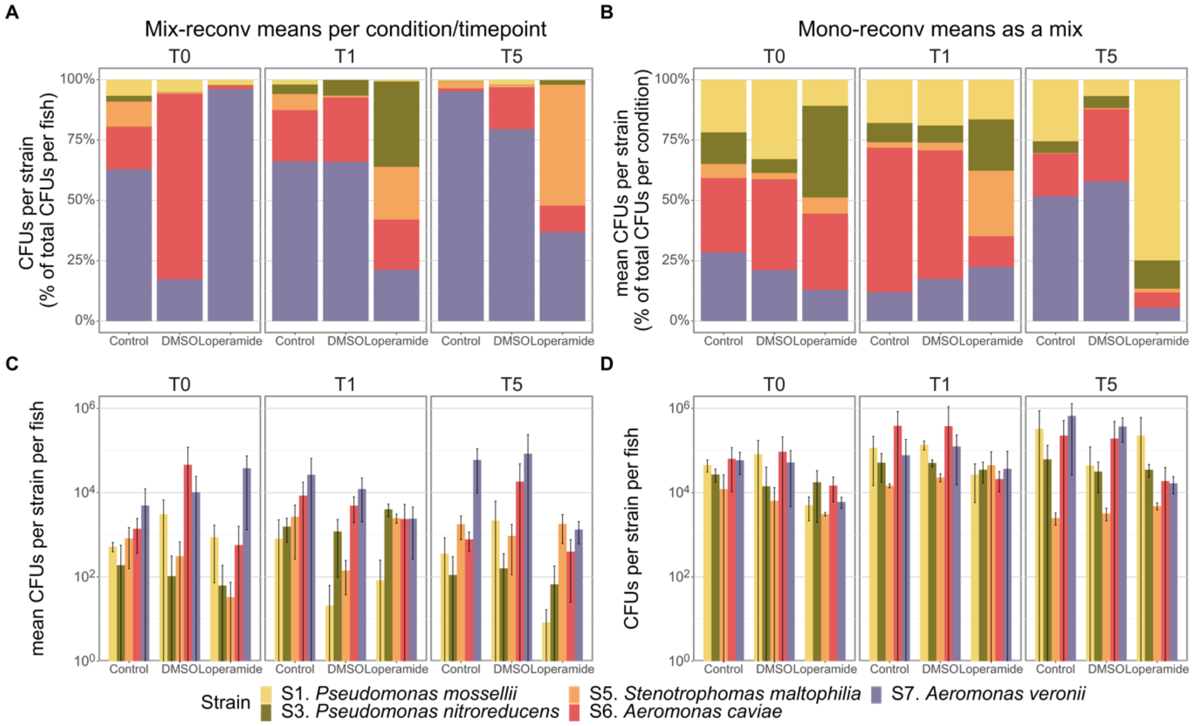


**Figure S9. Comparison of mono-reconventionalized colonization means with Mix-reconventionalized fish. (A)** Mix-reconventionalized means per condition and timepoint normalized to percent CFUs per fish. **(B)** Mono-reconventionalized fish colonization means combined as a hypothetical mix per condition and timepoint, normalized to percent CFUs per condition. **(C)** Mix-reconventionalized means per condition and timepoint as CFUs per fish per strain. **(D)** Mono-reconventionalized fish CFUs per fish per strain. For **C** and **D**: mean ± standard deviation per condition is shown on log scale (n = 3-4 fish).

**Table S1. Zebrafish environment bacterial strains used in this study**

| Code | Strain | Isolation Source | Reference |
|---|---|---|---|
| **S1** | *Pseudomonas mossellii* | Conventional zebrafish | [1] |
| **S2** | *Variovorax gossypii* | Conventional zebrafish | This study |
| **S3** | *Pseudomonas nitroreducens* | Conventional zebrafish | [1] |
| **S4** | *Achromobacter marplatensis* | Conventional zebrafish | This study |
| **S5** | *Stenotrophomas maltophilia* | Conventional zebrafish | [1] |
| **S6** | *Aeromonas caviae* | Conventional zebrafish | [1] |
| **S7** | *Aeromonas veronii* | Conventional zebrafish | [1] |
| **S8** | *Rhizobium sp.* | Conventional zebrafish | This study |
| **S9** | *Ochrobactrum tritici* | Conventional zebrafish | This study |
| **S10** | *Flavobacterium johnsoniae* | soil | [2] |

[1] F. A. Stressmann *et al.*, "Mining zebrafish microbiota reveals key community-level resistance against fish pathogen infection," *ISME Journal*, vol. 15, no. 3, pp. 702–719, 2021, doi: 10.1038/s41396-020-00807-8.

[2] R. A. Lewin and D. M. Lounsbery, "Isolation, cultivation and characterization of flexibacteria," *J Gen Microbiol*, vol. 58, no. 2, pp. 145–170, Oct. 1969, doi: 10.1099/00221287-58-2-145.
